# Rapid spread of nutritional *Wolbachia* symbionts across and within solitary bee species

**DOI:** 10.64898/2026.06.17.729582

**Authors:** Leopold Preuß, Marta Tischer, Anastasia Andrews, Panagiotis Theodorou, Christoph Bleidorn, Stefanos Siozios, Michael Gerth

**Affiliations:** Department of Zoology, Institute for Biology, Faculty of Natural Sciences I, Martin Luther University Halle-Wittenberg, Halle (Saale), Germany; German Centre for Integrative Biodiversity Research (iDiv) Halle-Leipzig-Jena, Leipzig, Germany; Institute of Evolutionary Biology, University of Warsaw, Warsaw, Poland; Department of Environmental Science, Radboud University Nijmegen, The Netherlands; Animal Evolution & Biodiversity, Georg-August-University Göttingen, Göttingen, Germany

## Abstract

Maternally inherited, intracellular Bacteria of the genus *Wolbachia* are extremely widespread among arthropods. Their evolutionary success is owed to frequent host shifts and rapid subsequent spread within host populations, often facilitated by *Wolbachia*-induced reproductive manipulations. Theory suggests that carrying *Wolbachia* must also be beneficial for a successful spread, however the nature of such benefits remains unclear.

Here, we demonstrate that *Wolbachia*’s success in many solitary bee species is strongly associated with the presence of a gene cassette enabling *Wolbachia* to synthesize Biotin (vitamin B7). This ability is absent from almost all other *Wolbachia* strains but common among bee associated strains. We show that *Wolbachia* occurs in up to 70% of all bee species and that strains carrying this rare genomic element have independently spread into numerous bee hosts recently. We further demonstrate that the presence of the biotin operon is associated with specialised diets in several bee species, suggesting a previously overlooked nutritional role of *Wolbachia* in this group of important pollinators. Overall, our results suggest that nutritional benefits provided by symbionts may represent a more widespread and important mechanism by which reproductive manipulators spread into new host species than previously appreciated.

## Introduction

The maternally inherited, intracellular endosymbiont *Wolbachia* (Alphaproteobacteria) is found in an estimated 50% of all arthropod species, making it the most common symbiont on earth^1,2^. This success is due to *Wolbachia*’s ability to establish in novel species via host shifts and spread within host populations through reproductive manipulation. In particular, cytoplasmic incompatibility (CI), the reduced fertility of *Wolbachia* free females when mating with males carrying *Wolbachia*, may lead to extraordinarily fast spread of *Wolbachia* within insect populations, as supported by mathematical modelling of ecological dynamics^3,4^, and observations from various host systems^3,5–7^. Likewise, the spread of *Wolbachia* into novel host species occurs frequently and rapidly^8,9^. However, models predict that upon initial transfer to new hosts, CI alone may not be sufficient to promote the spread of the symbiont^10,11^. As the frequency of *Wolbachia* positive males is initially low, matings with *Wolbachia-*free females are too rare to select for *Wolbachia* positive females. Only after *Wolbachia* frequency has passed an infection threshold in a host population, will it spread via CI, often to fixation. To explain *Wolbachia* spread before CI can have an effect, positive fitness effects of the symbiont, which may lower *Wolbachia*’s infection threshold must be considered^12,13^. Furthermore, some *Wolbachia* strains spread entirely without causing CI^5,14^, which may also be explained by fitness effects. Such benefits are thus expected to be crucial for *Wolbachia*’s spread within and across species, however evidence for positive *Wolbachia* effects in natural host populations is limited.

The best candidates for fitness benefits aiding in *Wolbachia* establishment are pathogen protection and nutritional supplementation^15^. *Wolbachia* can provide strong protection from RNA viruses^16^, and this phenotype may be transferred into novel hosts when *Wolbachia* shifts hosts^17,18^. Although already widely applied in biological control of human viruses through artificial *Wolbachia* transfer into vector species^19^, it is unclear if protection from pathogens is relevant for the establishment of *Wolbachia* in novel host species. *Wolbachia*’s effect on viruses in lab experiments is variable, showing no effects for some viruses, and enhancement of others^20–22^. Furthermore, surveys of natural arthropod populations and their viruses have revealed that viral blocking through *Wolbachia* may not be common^23–26^. Similarly, the evidence for *Wolbachia* increasing insect fitness by providing nutrients is mixed^27^. In *Drosophila*, laboratory experiments have demonstrated that the *Wolbachia* strain *w*Mel can increase fitness by buffering against several conditions of nutritional stress^28,29^, and possibly this phenotype underlies positive fecundity effects observed in several arthropod hosts^15^. However, *Wolbachia* may evolve to become beneficial in adaptation to novel hosts^30^, and it remains uncertain if such nutritional benefits are already present at the time of host shifts, when it would be required for *Wolbachia* to spread.

Unambiguous examples of positive fitness effects stem from filarial nematodes, in which *Wolbachia* is an obligate mutualist^31^. In insects, obligate mutualism is especially common in species feeding exclusively on plant sap or blood^32^. *Wolbachia* is one of many symbionts that may provide essential nutrients lacking in these diets, such as biotin or riboflavin^33,34^. The ability of *Wolbachia* to produce biotin (vitamin B7) is limited to a handful of strains^35^, and a gene cassette containing six biotin synthesis genes (biotin operon of obligate intracellular microbes – BOOM) is present in *Wolbachia* strains associated with blood and sap feeders^34,36,37^, as expected from their feeding ecology. However, BOOM was also found in a few *Wolbachia* strains of solitary bees^38^, a species rich group of ecologically and economically important pollinators^39,40^. *Wolbachia* occurs at high incidence across manny lineages within bees^41^, but little is known about the factors that shape the distribution and host shift patterns of *Wolbachia* in solitary bees^42^. Bees raise their offspring nearly exclusively on pollen, which is generally regarded as nutritious and rich in vitamins^43^. The puzzling presence of biotin producing *Wolbachia* symbionts in bees was thus hypothesized to be a rare and non-adaptive by-product of lateral genetic transfer between intracellular microbes^38^.

Here, by analysing more than 130 newly assembled complete and draft metagenomes together with available reference genomes, we show that the biotin operon is particularly widespread in *Wolbachia* symbionts within and across bee species, compared to other host taxa. Our data suggest that the successful spread of *Wolbachia* strains in different bee hosts coincided with, and is likely facilitated by, the spread of multiple BOOM operons across strains. We further show that the presence of BOOM is linked to the feeding ecology of specialized bees, hinting at a previously overlooked nutritional mutualism. Together, our findings are consistent with the hypothesis that nutritional benefits drive the spread of *Wolbachia* in new populations.

## Results and Discussion

### *Wolbachia* strains encoding biotin synthesis genes are generally rare, but very common in bees

We identified the BOOM in 83 out of 323 *Wolbachia* genomes analysed (26%, Supplementary Table 1) from hosts spanning seven arthropod orders. The vast majority of these 83 *Wolbachia* strains (76%) are associated with solitary bees, despite solitary bees representing only 26% of all investigated hosts. This highlights a strong enrichment of BOOM-containing *Wolbachia* in solitary bees compared to other arthropod hosts (chi^2^=42.032, p=8.98e-11). Such a strong taxonomic enrichment is very unlikely to arise by chance and argues for an important function of the operon in bee *Wolbachia*. BOOM was also found in hosts for which biotin complementation has been demonstrated or is very likely (blood or phloem feeders) and in four non-bee hymenopterans (Supplementary Table 1). In many cases, multiple *Wolbachia* infections (up to four) in a single bee host were identified, and in several instances (13 individuals from five bee species), multiple copies of the BOOM were found, either on a single or on separate *Wolbachia* genomes (Supplementary Table 1).

A targeted screen of 198 bee species revealed *Wolbachia* to be present in 141 species (71%) and the biotin operon in about half of the corresponding *Wolbachia* genomes (67/141) (Supplementary Figure 3). Further analysis of population genomic data for two bee species (*Andrena vaga* and *A. florea*) revealed that *Wolbachia* and its associated BOOM region were present in 100% of the individuals sampled across multiple populations (114 and 70, respectively), with almost no intraspecific genetic variation detected across the *Wolbachia* genomes, including the biotin operon (Supplementary Tables 2 & 3). The high degree of similarity in *Wolbachia* genomes across different host populations indicates a relatively recent origin and spread of *Wolbachia* in these species. For any other bee species for which multiple individuals could be investigated, we find that when *Wolbachia* and the BOOM are present, they are usually found in all individuals (Supplementary Table 4). This suggests *Wolbachia* and BOOM are very common and at fixation in several solitary bee species. The high incidence of *Wolbachia* within and across bee species is in line with the interpretation of BOOM carrying *Wolbachia* providing fitness benefits, through which they may facilitate their establishment and spread.

### Two origins of BOOM in bee *Wolbachia*, and many origins of *Wolbachia* in bees

A phylogenetic reconstruction of microbial biotin operons revealed that BOOM in *Wolbachia* are monophyletic with the exception of a single BOOM found in *Wolbachia* of the mite *Fragariocoptes setiger* (Fig. 1). Within the *Wolbachia* BOOM clade, BOOMs of bee-associated *Wolbachia* form two distinct evolutionary lineages (beeBOOM1 & beeBOOM2, Fig. 1), each with very little within-lineage sequence variation (beeBOOM1 99.1 - 100% and beeBOOM2 100% identity, estimated with fastANI^44^). This points to at least two independent evolutionary origins of BOOM in bee *Wolbachia*, both of which are likely recent.

**Figure 1.**
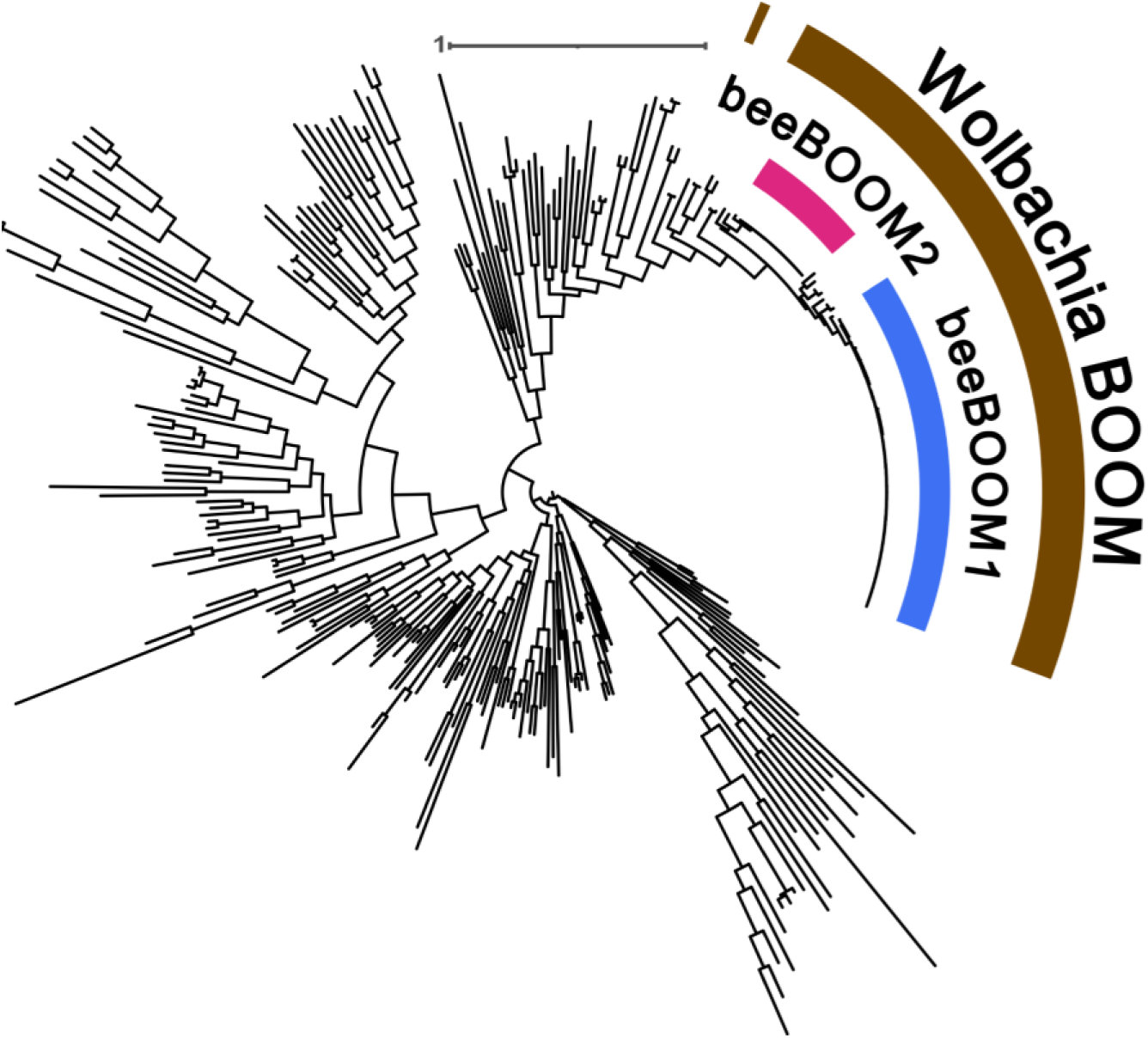
| Phylogeny of bacterial biotin operons. Midpoint rooted phylogeny of microbial biotin operons, inferred using IQ-Tree v3.1.1 under model Q.pfam+I+R10 as determined by Modelfinder Plus. *Wolbachia* BOOM is highlighted in brown, while the two BOOM clades of bee infecting *Wolbachia* strains are highlighted in blue and red. All leaf labels were omitted to increase visibility. Scale bar corresponds to the average number of inferred substitutions per site.

The phylogenetic reconstruction of 323 supergroup A and B *Wolbachia* genomes shows that the symbiont has spread into solitary bees multiple times independently and that BOOM is almost exclusively associated with supergroup A *Wolbachia* (Fig. 2). BeeBOOM1 is found in multiple phylogenetically distinct *Wolbachia* lineages, whereas beeBOOM2 is limited to a single, *w*Ri-like *Wolbachia* strain. *w*Ri-like strains are known for their rapid, recent spread into many different *Drosophila* species in which they lack a BOOM^8^. This means that either BOOM was ancestrally present in *w*Ri but was later lost multiple times, or BOOM was independently acquired by a single *w*Ri variant, which subsequently spread into many bee hosts. These two evolutionary scenarios cannot be confidently resolved because of the high degree of phylogenetic relatedness between the *w*Ri variants, reflecting the recent evolutionary history of these events.

**Figure 2.**
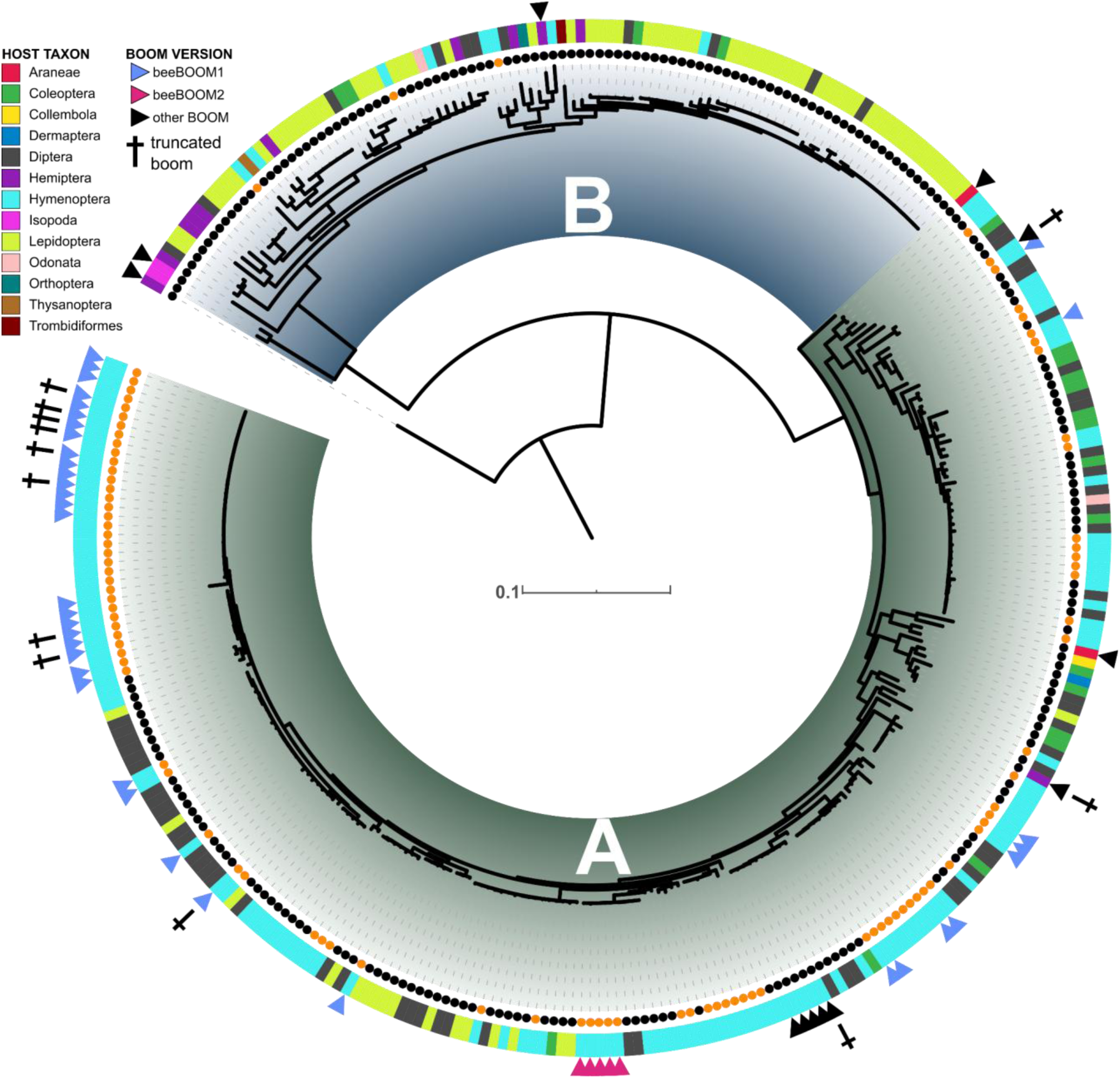
| Phylogeny of A and B supergroup *Wolbachia*. Phylogeny of A and B supergroup *Wolbachia* including all ‘reference genome’ level assemblies from NCBI as well as long and short read meta-assembled draft genomes from the current study. Reconstruction was done with IQ-Tree v3.1.1 under Q.BIRD+F+I+R5 model. Leaf labels were substituted with dots coloured by taxonomy: Orange indicates a host species belonging to solitary bees (Anthophila), Black corresponds to other taxa. The colours of the ring around the leaf labels indicate the arthropod order of the host species. Triangles pointing at individual leaves indicate the presence of BOOM on the respective *Wolbachia* genome, with blue and red triangles representing beeBOOM1 and beeBOOM2, respectively, while a black triangle indicates non-bee BOOM and the yellow triangle is assigned to the BOOM of a *Wolbachia* strain infecting *Augochloropsis metallica*, which did not cluster with the other two bee BOOM versions, likely due to heavy truncation. Truncated BOOMs are indicated with a cross symbol above the triangle.

In multiple instances, the same *Wolbachia* strain carrying BOOM is found in several unrelated bee hosts (Fig. 2). It is noteworthy that, in the few cases for which these *Wolbachia* strains were also found in non-bee hosts, they consistently lack the BOOM region.This pattern is consistent with BOOM loss following host shifts into other arthropod hosts. Our phylogenetic analyses are in line with a relatively rapid spread of certain *Wolbachia* lineages that carry BOOM into bees, but not other arthropod species. Furthermore, the high number of distinct bee-associated *Wolbachia* lineages carrying related BOOMs strongly suggests that the operon is regularly acquired by *Wolbachia* horizontally. While we do not know the rate by which BOOM is introduced into different *Wolbachia* strains, our data suggest that the operon is only retained in strains associated with bees, consistent with a functional role in these hosts.

### Evolution of beeBOOM is driven by prophage dynamics

Most BOOM operons of bee *Wolbachia* were located within predicted phage regions or in close genomic proximity to phage genes (Supplementary Figure 1 and Supplementary Table 5). Taxonomically, these phages were classified as phage WO (Caudoviricetes), which is commonly found in *Wolbachia* and often the carrier for functionally important loci (e.g., the loci responsible for CI and male killing)^45^. The association of the BOOM with these phage elements suggests it also facilitates the introduction and the dissemination of the operon between bee infecting *Wolbachia* strains. Phylogenetic analyses of BOOM regions and associated phages show that the two variants of beeBOOM associate with different phage lineages (Fig. 3). The phages associated with beeBOOM1 are not monophyletic and cluster with phages of non-bee BOOMs (Fig. 3). In some cases, beeBOOM1 was associated with phages showing clear signs of deterioration and genome rearrangements (Fig. 4; Supplementary Figure 2). Deterioration of other phage loci compared to the BOOM operons is in line with purifying selection acting on BOOM, arguing for its functional role in bee-associated *Wolbachia*. However, truncation and loss of beeBOOM1 are also occasionally observed in bee *Wolbachia*, driven by activity of transposases adjacent to the BOOM region (Fig. 4A). The beneficial effect of biotin provisioning may thus be only required for *Wolbachia* during the initial establishment in bee species after host shifts. As predicted by modelling^12^, high *Wolbachia* incidence rates may be maintained by CI alone once a threshold frequency has been reached, potentially rendering the BOOM obsolete in some cases.

**Figure 3.**
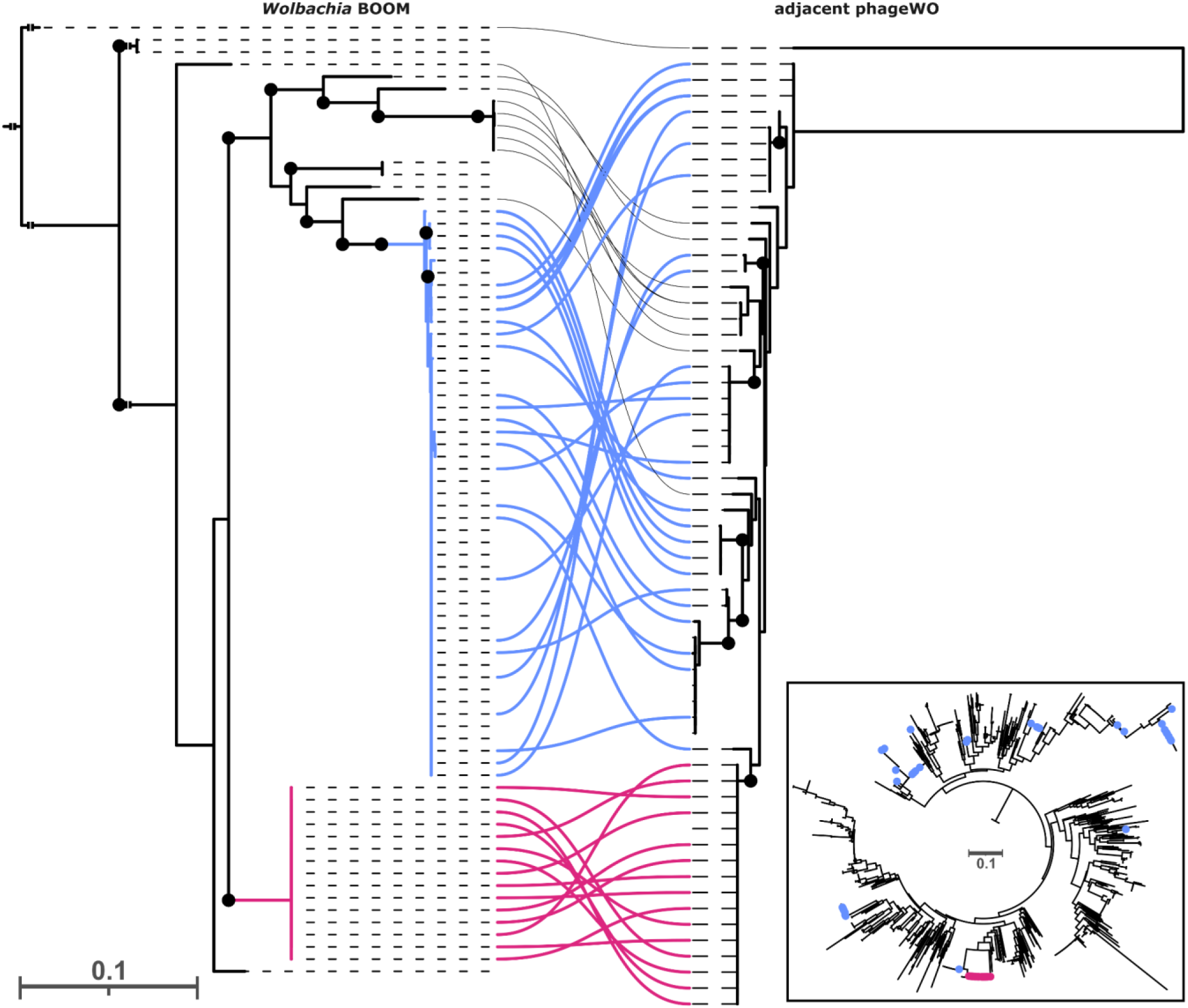
| Cophylogeny of *Wolbachia* BOOM and associated phages. Phylogenetic reconstruction of beeBOOM (left) with BOOM of *Candidatus* Mesenet endosymbiont of *Agriotes lineatus* as outgroup. Ultrafast bootstrap values >98 are indicated with a black circle on the nodes. The phage phylogeny (right) is based on the large serine recombinase gene of phage WO if it was found in proximity to the BOOM on the contig. The inset shows the phylogeny of all PhageWO serine recombinase loci in the dataset, including from non-BOOM-carrying and non-bee-infecting *Wolbachia* strains. Both phylogenies were constructed using IQ-Tree v3.1.1 with model TVM+F+I+R2 for BOOM and model JTT+F+R4 for the phage locus. Blue and red colours represent beeBOOM1 and beeBOOM2, respectively and lines connect BOOM with the associated phage on the same genome. Scale bar corresponds to estimated average number of substitutions per site.

**Figure 4.**
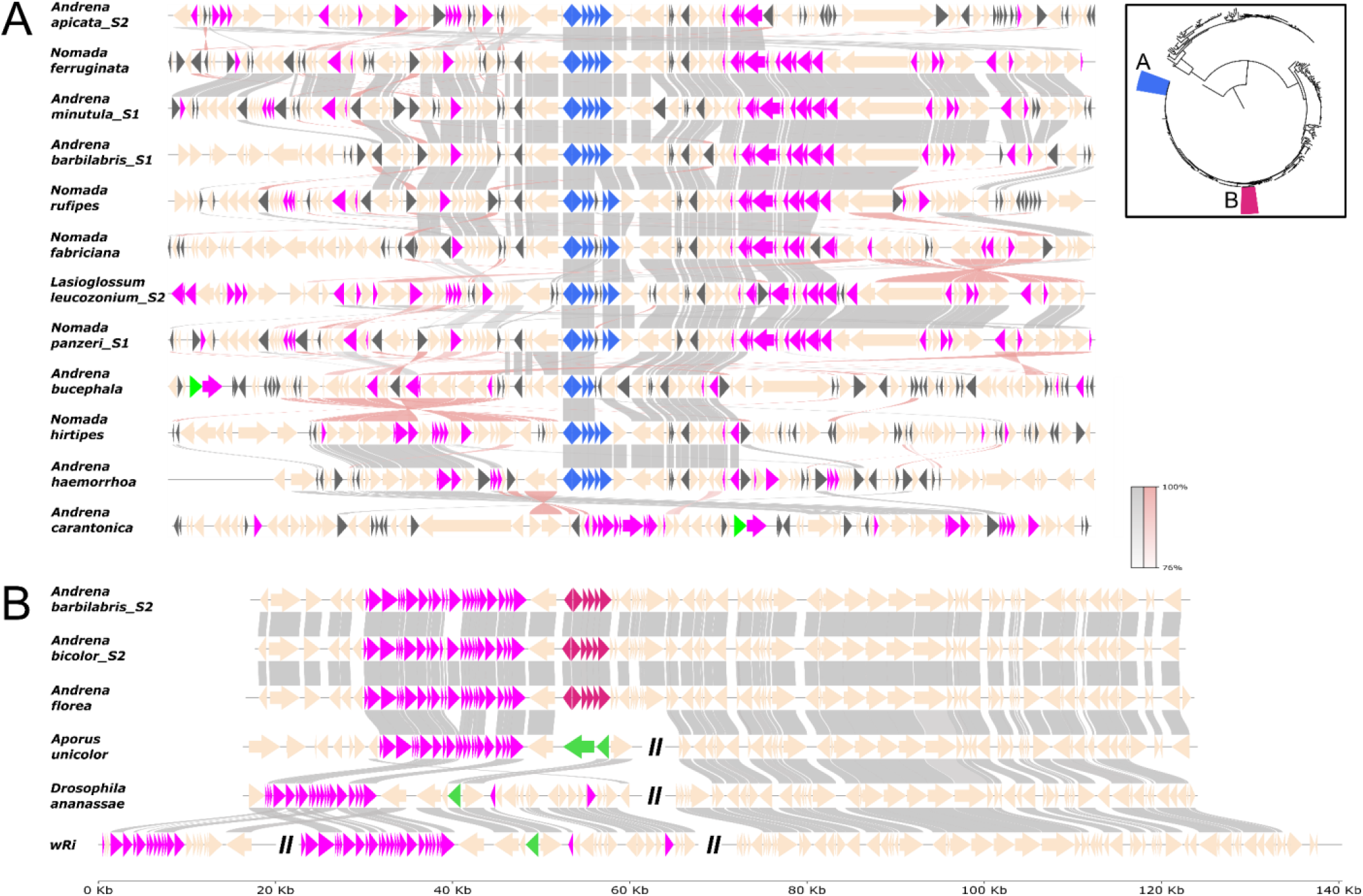
| Genomic regions surrounding beeBOOMs. Synteny plots of beeBOOM1 (**A**) and beeBOOM2 (**B**) showing features ∼ 50k bp up- and downstream of the biotin operon. Grey links indicate gene similarity >70% assessed by MMseqs. Blue and red arrows represent the genes comprising beeBOOM1 and beeBOOM2, respectively. Pink arrows indicate phage-associated genes, green arrows cifA/cifB genes and black arrows represent transposases. For non-BOOM carrying genomes from the *w*Ri clade, multiple regions were included to highlight the genomic rearrangements in comparison with the other strains. A double line indicates a visualisation break in the genome.

In contrast, beeBOOM2 region is associated with a monophyletic clade of phages and appears to be intact (Figs. 3 & 4). The phages associated with beeBOOM2 lack genetic differentiation, and are only found in a single, *w*Ri-like *Wolbachia* lineage (Figs. 2 & 3). This further supports a recent, phage-mediated origin of beeBOOM2 in one *Wolbachia* strain that subsequently spread into several bee hosts. The phage regions surrounding beeBOOM2 are syntenic and the genomes of the corresponding *Wolbachia* strains highly similar (99.7% average nucleotide identity, Supplementary Table 6). These observations, together with pseudogenization of phage and BOOM loci in beeBOOM1, but not beeBOOM2, argue for a more recent origin of beeBOOM2 compared with beeBOOM1. Interestingly, *Wolbachia* carrying beeBOOM2 preferentially spread into bee hosts already carrying a *Wolbachia* strain with the other version of the operon. In these cases, the BOOM may be functionally redundant, but apparently still provides sufficient benefits for the host in order to help *Wolbachia* establish. Theoretical models predict that the infection threshold for less competitive strains may be lowered by the presence of coinfecting strains^46^, so the *Wolbachia* strain carrying beeBOOM2 may benefit from the presence of the strain carrying the other BOOM variant. Alternatively, this could indicate an ongoing replacement of one BOOM carrying *Wolbachia* strain with another, reminiscent of a recent *Wolbachia* replacement observed in *Drosophila melanogaster*^47^.

### The presence of *Wolbachia* strains carrying bee BOOM is linked to pollen preference of specialised bee hosts

The presence of the biotin operon in genomes of vertically inherited symbionts of arthropods usually points towards a nutritional symbiosis in which the symbiont augments specialised host diets (often blood or phloem) with biotin^48^. Nutritional specialisation also occurs in solitary bees, as many species use only a single or very few plant species as pollen sources^49^. Based on a literature survey of bee ecology^55^, we find that among the bee species that carry *Wolbachia* strains with biotin operons, specialised bees are not overrepresented (Fig. 5A). However, indicator species analyses revealed that a nutritional specialisation on willow (*Salix* sp.) pollen is significantly associated with the presence of *Wolbachia* carrying a biotin operon (p=0.002). Indeed, all six bee species in our dataset that feed exclusively on willow pollen harbour *Wolbachia* strains carrying BOOM. This is also true for bees specialised on bryony (*Bryonia* sp.), or the closely related widow flowers and scabious (*Knautia* sp. and *Scabiosa* sp.), although the small sample number precludes statistical validation in these cases. Our observations suggest that biotin supplementation is not strictly required for most bee species, but may be necessary for some highly specialised bees with a very uniform diet that potentially does not contain sufficient amounts of biotin.

**Figure 5.**
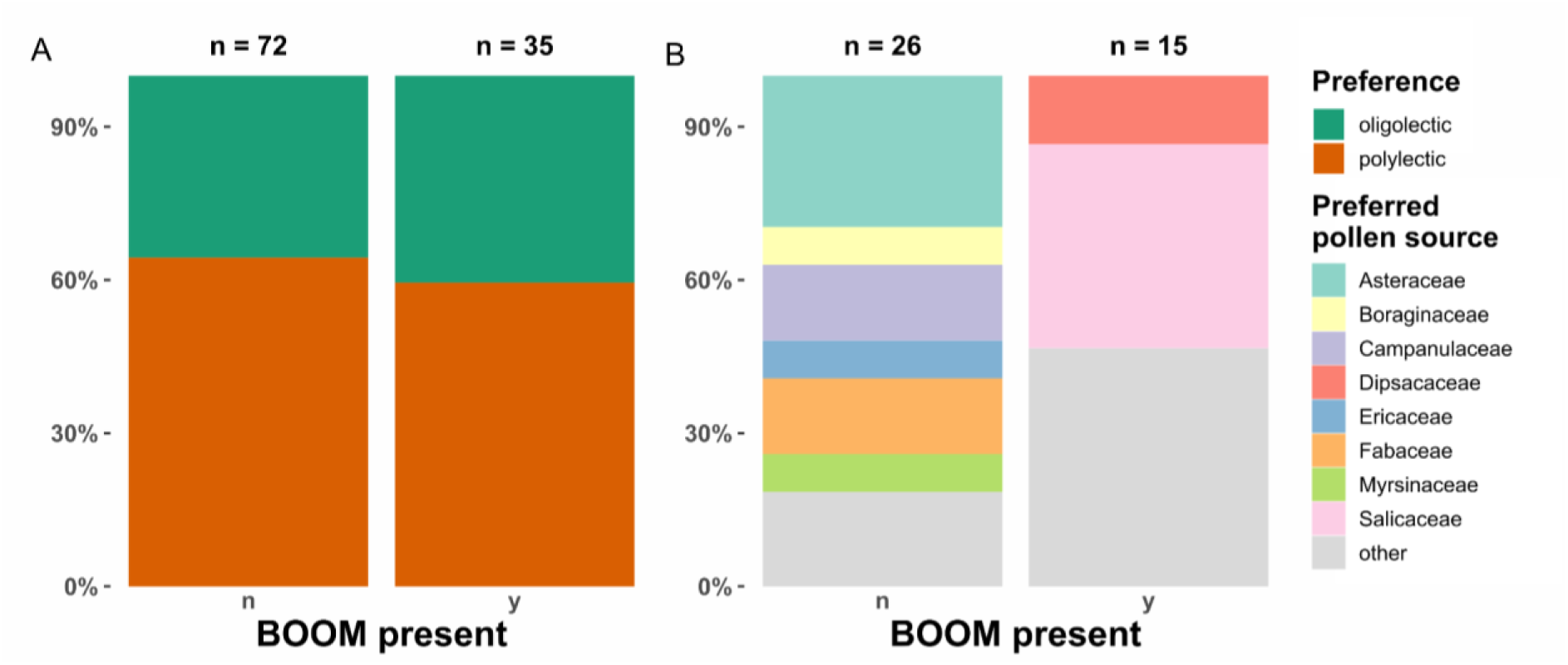
| Feeding ecology of bees with and without BOOM carrying *Wolbachia*. **A**: Degree of specialisation of solitary bee species on specific host plant taxa. Only pollen collecting bees were included Polylectic: collecting pollen from a broad spectrum of host plants, Oligolectic: collecting pollen from a single or very few related host plant species according to literature data^55^. **B**: Preferred pollen source of oligolectic solitary bee species with and without BOOM. Different colours represent different plant families. Families that are visited by less than 7% of bee species in each group were summarised as ‘other’.

Although pollen is generally regarded as nutritious^50^, very little comparative data exist on the nutritional value of pollen produced by individual plant species^51^, including those required by highly specialised bees. Conceivably, there is natural variation in the vitamin content of pollen from different plant species. Due to the experimentally intractable nature of the ground nesting solitary bee lifestyle, the interpretation of the physiological role and importance of biotin supplementation in solitary bees remains speculative. Bee pollen is predominantly used as provision for developing larvae^49^, and biotin is a cofactor in various metabolic reactions^52^. Biotin supplied by *Wolbachia* could thus directly benefit bee growth and development. Alternatively, biotin could be important to select for beneficial microbes in the nest or brood capsule, which may make the pollen accessible through detoxification and fermentation^53^ or even serve directly as a food source for the developing larvae^54^. The wide distribution of BOOM in *Wolbachia* symbionts of bees suggests that the benefit of biotin provisioning is not limited to specialist bees, and may be generally important for *Wolbachia* during initial stages of establishment in new bee hosts.

## Conclusion

*Wolbachia* symbionts are known for their ability to rapidly spread into many different host species. So far, this success has largely been attributed to *Wolbachia*’s manipulation of host reproduction, although symbiont mediated fitness benefits have long been hypothesised to facilitate the initial establishment of *Wolbachia* in novel host species. Here, we show that the capacity for biotin supplementation, previously considered rare and restricted to a small number of specialised *Wolbachia* strains, is widespread among *Wolbachia* symbionts of bees and is strongly associated with their repeated spread across bee hosts. Our findings suggest that nutritional provisioning by *Wolbachia* may be more common than previously appreciated and could represent an important factor contributing to the evolutionary success of these symbionts in bees.

## Methods

### Arthropod sampling and identification

Arthropod sampling was conducted between 2019 and 2021 around Göttingen city (Lower Saxony, Germany) during flowering season between March and September on four different host plant species *Medicago sativa* (Fabaceae), *Daucus carota* (Apiaceae), *Salix spp.* (Salicaceae, *Salix caprea* most dominant) and *Urtica dioica* (Urticaceae), which were identified on site if possible. The specimens were collected using an entomological exhauster with an attached Falcon™ tube and either frozen at −20 °C or preserved in 70-98 % ethanol. Arthropods were morphologically identified to the highest possible taxonomic level and grouped into morphospecies where species identification was not possible. Additionally, specimens for which species identification was missing were identified using the molecular barcoding marker COI (for details see processing of read data).

### DNA extraction and sequencing of insect pollinators

Generally, larger insects were pinned and dried and DNA extracted from thorax or leg musculature while DNA of smaller insects was extracted using the whole body. Microscopically small insects were pooled by morphospecies for DNA extraction. The tissue was frozen at −80 °C before homogenisation using DNA-free sterile plastic pestles in Eppendorf Tubes. All samples were incubated with proteinase K at 56 °C overnight and then processed with three different kits depending on initial insect size and host species. Small and fragile individuals were processed using the Qiagen DNeasy Blood & Tissue Kit, while the remaining samples were processed using Zymo Research Quick-DNA Kits and EurX Tissue & Bacterial DNA Purification Kit following the provided company protocols. *Wolbachia* infection was confirmed using PCR of conserved bacterial housekeeping genes (*coxA, dnaA, gatB, fbpA, ftsZ, hcpA, wsp*)^56,57^.

Altogether, 126 DNA extracts of *Wolbachia* infected samples (Supplementary Table 7) were used for genomic library preparation for low-coverage sequencing using Illumina technology. DNA at a concentration of 10 µg/µl was sheared using the sonicator Diagenode Bioruptor® Pico with 13 cycles of 30sec on/off sonication to obtain DNA fragments with 200-500 bp size. Fragmented DNA was processed stepwise as blunt-end repair where 5’- and 3’-ends are filled in or removed by T4 DNA polymerase (T4 DNA Polymerase (New England Biolabs, Ipswich, MA, USA; Cat. No. M0203), 5-phosphates are attached using T4 polynucleotide kinase (New England Biolabs, Ipswich, MA, USA; Cat. No. M0201). Illumina-compatible P5 and P7 adapter oligonucleotides (adapter sequences based on the standard Illumina P5 and P7 flow-cell binding sequences) were custom-synthesized by Eurofins Genomics (Ebersberg, Germany) are ligated to both ends with the T4 DNA ligase (New England Biolabs, Ipswich, MA, USA; Cat. No. M0202). Samples were indexed with two unique indexes using AccuPrime™ Taq DNA Polymerase High Fidelity (Invitrogen, Thermo Fisher Scientific, Waltham, MA, USA; Cat. No. 12346-086). Additional PCR reactions with Illumina-compatible indexed adapters IS5 and IS6 (adapter design was based on standard Illumina adapter architecture, custom synthesized by Eurofins Genomics, Ebersberg, Germany), were carried out to avoid indexing reaction- heteroduplex constructs^58,59^. This process was finalised using the MinElute PCR Purification Kit (QIAGEN, Hilden, Germany; Cat. No. 28004) for cleanup. During library prep cleanup and size-selection was done using Agencourt AMPure XP magnetic beads (Beckman Coulter, Brea, CA, USA; Cat. No. A63880). Each step included quality control with the Qiaxcel OM00-OL00 method (QIAxcel Advanced System (QIAGEN, Hilden, Germany). Finally, six pools representing 126 individuals were sequenced by Novogene, Advancing Genomics Facility in the United Kingdom.

### Processing of metagenomic sequencing data

Raw Illumina reads for each sample were merged after adapter trimming and quality control with fastp^60^ (v0.23.0) and subsequently meta-assembled with megahit^61^ (v1.2.9). The meta-assemblies were filtered and binned using BlobToolKit^62^ (v4.4.4) which required coverage information and BLAST hits. Coverage files were created using minimap2^63^(v2.28-r1209). BLAST hits were generated by performing a BLASTn search against a local copy of the NCBI prokaryotic RefSeq database (downloaded March 2nd 2025) using the BLAST command line application^64^ (v2.16.0+) with settings as recommended in BlobToolKit documentation (-outfmt 6 -max_target_seqs 10 -max_hsps 1 -evalue 1e-25). Resulting bins were then filtered for the genus *Wolbachia* and a minimum size of 1Mb, and quality assessed using CheckM^65^(v1.2.4). Bins for which size and CheckM results indicated the presence of two separate *Wolbachia* strains, which could not be properly separated, were excluded from the *Wolbachia* strain phylogeny as these double infections drastically reduce the amount of single copy orthologues present and introduce artifacts in the phylogeny. If BlobToolKit yielded multiple bins for the same sample, these were separately processed as different strains belonging to the same host species.

In addition to the Illumina reads of *Wolbachia* infected arthropods, unassembled PacBio sequencing reads for solitary bees were obtained from the NCBI Sequencing Read Archive (SRA). For a detailed list of samples and their origin see Supplementary Table 1. The raw reads were mapped with minimap2 against a local reference database containing all available reference genomes for *Wolbachia* on NCBI. Fasta sequences were extracted for perfectly matched reads using SAMtools^66^ (v1.22.1) and then assembled with the Flye assembler^67^ (v2.9.5-b1801). The resulting assemblies were inspected and exported to fasta sequences using the interactive assembly viewer Bandage^68^ (v.0.8.1), whereby each assembly bubble between 1-2mbp of size was considered a separate, true *Wolbachia* genome.

### *Wolbachia* phylogeny reconstruction

The *Wolbachia* strain phylogeny was created using a pipeline based on Snakemake^69^ (v.9.5.1.). In short, publicly available reference genomes for the genus *Wolbachia* were downloaded by NCBI datasets^70^ (v16.40.1) and annotated together with additional genomes provided to the pipeline using Bakta^71^ (v1.11.4). Single copy orthologues were identified and extracted by SonicParanoid2^72^ (v2.0.8) before alignment construction with MAFFT^73^ (v7.525), resulting in 98 single copy ortholog alignments. IQ-Tree2^74^ (v2.4.0) was then used for concatenation of alignments, model selection with ModelFinder plus, tree reconstruction and ultrafast bootstrap estimation with 1000 pseudoreplicates. The bedbug infecting *Wolbachia* strain *wCle* (supergroup F) served as an outgroup for the strain phylogeny.

### PCR screen for BOOM carrying *Wolbachia* in solitary bees

To estimate the frequency of BOOM carrying *Wolbachia* infection in solitary bees, samples of extracted DNA representing 174 different bee species collected in Germany^75^ were used for a PCR screen. Wolbachia infection was detected through amplification of *ftsZ* primers specific for *Wolbachia*^76^ and presence of BOOM was inferred using primers specific for the bioA gene^38^. PCR results were inspected on agarose gel using Mupid One gel system. Amplified bioA fragments were further sequenced by Eurofins Genomics to confirm specificity of the PCR. Primer sequences and PCR program details can be found in the Supplementary Table 8. The PCR screen was complemented by genomic data for cases in which no DNA extract was available for PCR.

In some cases bioA could not be detected via PCR screen while it was present in the metagenomic sequence data from other individuals. These species were regarded as both *Wolbachia* and beeBOOM positive as the age and possibly reduced quality of DNA samples used for PCR screen likely explain failed detection of BOOM using this method.

### BOOM and phage detection and phylogeny reconstruction

Presence of BOOM in the *Wolbachia* strains was detected by blasting the protein sequences of each predicted CDS against a local reference database consisting of the BOOM of the *Wolbachia* strain infecting *Cimex lectularius* (*wCle*). The BLAST results were filtered for percent identity and length as follows: bioA 60% and 350AA, bioB 65% and 270AA, bioC 60% and 170AA, bioD 65% and 190AA, bioF 60% and 300AA, bioH 60% and 200AA. For samples with hypothesized inseparable double infections, BOOMs were only included and handled separately if they assembled on separate contigs. Also, only operons with at least five of the BOOM genes present after filtering were considered in the phylogenetic reconstruction. If one or more genes of the operon were absent from high quality, chromosome level assemblies the operons were considered truncated. Furthermore, when BLAST results indicated potential truncation of one or multiple genes of the operon, alignments were inspected manually. If a premature stop codon could be found or the operon was interrupted by a transposase, the gene, and consequently the entire operon, was considered truncated.

For all samples, geNomad^77^ (v1.11.0) was used for prophage region prediction, phage gene annotation and taxonomic classification. As suggested by Bordenstein & Bordenstein (2022)^45^ the viral gene for the large serine recombinase was used for reconstruction of the *Wolbachia* bacteriophage WO. In a first, conservative approach, the protein sequence for this gene was taken from the geNomad annotations of all samples to ensure that the recombinase was in fact part of a predicted prophage which still retained all necessary genes for reentering the lytic phase. A second, less strict approach consisted of performing a local BLAST search of all orthogroups inferred by SonicParanoid against a reference serine recombinase sequence (Genbank QBB84083.1) to identify orthogroups representing this gene. The orthogroups were filtered for genes with a length between 200 and 600 amino acids which were then subjected to phylogenetic analysis. As described for the *Wolbachia* strain phylogeny, MAFFT was used for sequence alignment and IQ-Tree2 for alignment concatenation, model selection, phylogenetic reconstruction and ultrafast bootstrap estimation. However, the beeBOOM phylogeny is based on nucleotides instead of amino acid alignments due to the high degree of similarity between operons. All phylogenies were visualised using iTOL. For synteny plots, genomes were reoriented using DNAapler (v1.2.0) and genome visualisation was done with pyGenomeViz (v1.6.1) and R^78^ in the R studio environment^79^. The phylogeny containing all microbial BOOMs was reconstructed based on an alignment kindly provided by Michele Castelli, PhD and Leandro Gammuto, PhD, based on their recently published study (Giovanni et al., 2026^80^).

### Statistical analysis of pollen preference and BOOM occurrence

Pollen preferences and host plant spectrum of bee species were taken from Westrich (2019)^55^. We tested for host plants with statistically significant association with beeBOOM presence through indicator species analysis on a Bray-Curtis distance matrix using the indval function of the R package vegan^81^. Plots on pollen preference were created using the package ggplot2 in the R studio environment.

### Population genomics of *Wolbachia* endosymbionts of *Andrena vaga* and *Andrena florea*

Genomes of *Wolbachia* strains infecting the two mining bee species *Andrena vaga* and *Andrena florea* were assembled from PacBio long-reads generated by Baltz et al. (2026)^82^. as described above for *Wolbachia* genome assembly from unassembled PacBio reads. This resulted in two separate *Wolbachia* genomes from the *Andrena vaga* sample, one of which was circular, and one from the *Andrena florea* sample. These were subsequently used as reference for population wide SNP calling. Population level illumina reads for *A. vaga* and *A. florea* were obtained from Baltz et al. (2026)^82^. Quality control for each sample was done using fastp prior to SNP calling with snippy v4.6.0 under default options and core SNP summary with snippy –core.

## Acknowledgements

LP and MG are funded by the Deutsche Forschungsgemeinschaft (DFG, German Research Foundation) – 497854142. CB and MT were funded by the DFG – 414708180. The work was supported by the German Centre for Integrative Biodiversity Research (iDiv), funded by the DFG (FZT 118, 202548816).

## Supplementary data

**Supplementary Figure 1 | Genomic position of prophageWO regions BOOM and large serine-recombinase phage locus in investigated *Wolbachia* GENOMES found in bee infecting strains.** Phage regions were predicted with GENOMAD and arrows indicate loci used for phage phylogeny reconstruction shown in figure 3. The length of the grey bar indicates the length of the investigated contig. Only contigs > 600kB were considered for this figure.

**Supplementary Figure 2 | Full genome synteny for the *Wolbachia* genome excerpts shown in Figure 4.** A and B correspond to samples taken from two different lineages of *Wolbachia* highlighted in the inset.

**Supplementary Figure 3 | Summarised result of targeted screen for *Wolbachia* and BOOM in solitary bees combining PCR and metagenomic results.**

**Supplementary Table 1 | Main sample overview table.** Table of samples including Genbank accessions for reference genomes, SRA accessions for newly meta-assembled genomes from the SRA database, assembly stats, checkM results, host information and presence of boom, riboflavin and cif genes

**Supplementary Table 2 | Core SNPS for *Andrena vaga* population samples.** Core SNPS were identified using snippy-core.

**Supplementary Table 3 | Core SNPS for *Andrena florea* population samples.** Core SNPS were identified using snippy-core.

**Supplementary Table 4 | BOOM and *Wolbachia* screening overview.** List and results of bee species screened for the presence of *Wolbachia* and BOOM via PCR and/or metagenomic screen.

**Supplementary Table 5 | Genome coordinates for BOOM and adjacent prophageWO regions,including virus topology, length and number of genes.** BOOM gene positions were identified through BLAST as described in the methods and then summarized to coordinates for the entire BOOM. Prophage WO regions were predicted by running GENOMAD on all BOOM containing contigs.

**Supplementary Table 6 | Average nucleotide similarity of *Wolbachia* draft- and reference genomes used in this study estimated with fastANI.** X1 & X2: Compared Isolates; X3: Estimated ANI; X4: Orthologous matches; X5: Total number of sequence fragments.

**Supplementary Table 7 | Sample overview of Illumina sequenced pollinator libraries.**

**Supplementary Table 8 | Primer sequences and PCR programs used for targeted *Wolbachia* and BOOM screen in solitary bees**

## Literature

Weinert, L.A., Araujo-Jnr, E.V., Ahmed, M.Z., and Welch, J.J. (2015). The incidence of bacterial endosymbionts in terrestrial arthropods. Proc. Biol. Sci. 282, 20150249. 10.1098/rspb.2015.0249.

Hilgenboecker, K., Hammerstein, P., Schlattmann, P., Telschow, A., and Werren, J.H. (2008). How many species are infected with Wolbachia?--A statistical analysis of current data. FEMS Microbiol. Lett. 281, 215–220. 10.1111/j.1574-6968.2008.01110.x.

Turelli, M., and Hoffmann, A.A. (1991). Rapid spread of an inherited incompatibility factor in California Drosophila. Nature 353, 440–442. 10.1038/353440a0.

Engelstädter, J., and Telschow, A. (2009). Cytoplasmic incompatibility and host population structure. Heredity 103, 196–207. 10.1038/hdy.2009.53.

Kriesner, P., Hoffmann, A.A., Lee, S.F., Turelli, M., and Weeks, A.R. (2013). Rapid Sequential Spread of Two Wolbachia Variants in Drosophila simulans. PLOS Pathog. 9, e1003607. 10.1371/journal.ppat.1003607.

Jiggins, F.M. (2017). The spread of Wolbachia through mosquito populations. PLoS Biol. 15, e2002780. 10.1371/journal.pbio.2002780.

Bakovic, V., Schebeck, M., Telschow, A., Stauffer, C., and Schuler, H. (2018). Spatial spread of Wolbachia in Rhagoletis cerasi populations. Biol. Lett. 14, 20180161. 10.1098/rsbl.2018.0161.

Turelli, M., Cooper, B.S., Richardson, K.M., Ginsberg, P.S., Peckenpaugh, B., Antelope, C.X., Kim, K.J., May, M.R., Abrieux, A., Wilson, D.A., et al. (2018). Rapid Global Spread of wRi-like Wolbachia across Multiple Drosophila. Curr. Biol. 28, 963–971.e8. 10.1016/j.cub.2018.02.015.

Sanaei, E., Charlat, S., and Engelstädter, J. (2021). Wolbachia host shifts: routes, mechanisms, constraints and evolutionary consequences. Biol. Rev. 96, 433–453. 10.1111/brv.12663.

Fine, P.E.M. (1978). On the dynamics of symbiote-dependent cytoplasmic incompatibility in culicine mosquitoes. J. Invertebr. Pathol. 10.1016/0022-2011(78)90102-7.

Turelli, M., and Barton, N.H. (2017). Deploying dengue-suppressing Wolbachia : Robust models predict slow but effective spatial spread in Aedes aegypti. Theor. Popul. Biol. 10.1016/j.tpb.2017.03.003.

Fenton, A., Johnson, K.N., Brownlie, J.C., and Hurst, G.D.D. (2011). Solving the Wolbachia paradox: modeling the tripartite interaction between host, Wolbachia, and a natural enemy. Am. Nat. 178, 333–342. 10.1086/661247.

Zug, R., and Hammerstein, P. (2018). Evolution of reproductive parasites with direct fitness benefits. Heredity 120, 266–281. 10.1038/s41437-017-0022-5.

Kriesner, P., and Hoffmann, A.A. (2018). Rapid spread of a Wolbachia infection that does not affect host reproduction in Drosophila simulans cage populations. Evolution 72, 1475–1487. 10.1111/evo.13506.

Zug, R., and Hammerstein, P. (2015). Bad guys turned nice? A critical assessment of Wolbachia mutualisms in arthropod hosts. Biol. Rev. 90, 89–111. 10.1111/brv.12098.

Teixeira, L., Ferreira, Á., and Ashburner, M. (2008). The Bacterial Symbiont Wolbachia Induces Resistance to RNA Viral Infections in Drosophila melanogaster. PLOS Biol. 6, e1000002. 10.1371/journal.pbio.1000002.

Walker, T., Johnson, P.H., Moreira, L.A., Iturbe-Ormaetxe, I., Frentiu, F.D., McMeniman, C.J., Leong, Y.S., Dong, Y., Axford, J., Kriesner, P., et al. (2011). The wMel Wolbachia strain blocks dengue and invades caged Aedes aegypti populations. Nature 476, 450–453. 10.1038/nature10355.

Ant, T.H., Herd, C.S., Geoghegan, V., Hoffmann, A.A., and Sinkins, S.P. (2018). The Wolbachia strain wAu provides highly efficient virus transmission blocking in Aedes aegypti. PLOS Pathog. 14, e1006815. 10.1371/journal.ppat.1006815.

Marinotti, O., Paldi, N., and Gorla, D. (2026). Wolbachia releases for dengue control: why the evidence supports a pause and independent re-evaluation. Infect. Dis. 0, 1–11. 10.1080/23744235.2026.2668001.

Hussain, M., Lu, G., Torres, S., Edmonds, J.H., Kay, B.H., Khromykh, A.A., and Asgari, S. (2013). Effect of Wolbachia on Replication of West Nile Virus in a Mosquito Cell Line and Adult Mosquitoes. J. Virol. 87, 851–858. 10.1128/jvi.01837-12.

Dodson, B.L., Pujhari, S., Brustolin, M., Metz, H.C., and Rasgon, J.L. (2024). Variable effects of transient Wolbachia infections on alphaviruses in Aedes aegypti. PLoS Negl. Trop. Dis. 18, e0012633. 10.1371/journal.pntd.0012633.

Pimentel, A.C., Cesar, C.S., Martins, A.H.B., Martins, M., and Cogni, R. (2025). Wolbachia Offers Protection Against Two Common Natural Viruses of Drosophila. Microb. Ecol. 88, 24. 10.1007/s00248-025-02518-z.

Webster, C.L., Waldron, F.M., Robertson, S., Crowson, D., Ferrari, G., Quintana, J.F., Brouqui, J.-M., Bayne, E.H., Longdon, B., Buck, A.H., et al. (2015). The Discovery, Distribution, and Evolution of Viruses Associated with Drosophila melanogaster. PLOS Biol. 13, e1002210. 10.1371/journal.pbio.1002210.

Altinli, M., Lequime, S., Atyame, C., Justy, F., Weill, M., and Sicard, M. (2020). Wolbachia modulates prevalence and viral load of Culex pipiens densoviruses in natural populations. Mol. Ecol. 29, 4000–4013. 10.1111/mec.15609.

Ortiz-Baez, A.S., Shi, M., Hoffmann, A.A., and Holmes, E.C. (2021). RNA virome diversity and Wolbachia infection in individual Drosophila simulans flies. Preprint at bioRxiv, 10.1101/2021.05.09.443333 https://doi.org/10.1101/2021.05.09.443333.

Cogni, R., Ding, S.D., Pimentel, A.C., Day, J.P., and Jiggins, F.M. (2021). Wolbachia reduces virus infection in a natural population of Drosophila. Commun. Biol. 4, 1327. 10.1038/s42003-021-02838-z.

Newton, I.L.G., and Rice, D.W. (2020). The Jekyll and Hyde Symbiont: Could Wolbachia Be a Nutritional Mutualist? J. Bacteriol. 202, 10.1128/jb.00589-19. https://doi.org/10.1128/jb.00589-19.

Brownlie, J.C., Cass, B.N., Riegler, M., Witsenburg, J.J., Iturbe-Ormaetxe, I., McGraw, E.A., and O’Neill, S.L. (2009). Evidence for Metabolic Provisioning by a Common Invertebrate Endosymbiont, Wolbachia pipientis, during Periods of Nutritional Stress. PLOS Pathog. 5, e1000368. 10.1371/journal.ppat.1000368.

Lindsey, A.R.I., Tennessen, J.M., Gelaw, M.A., Jones, M.W., Parish, A.J., Newton, I.L.G., Nemkov, T., D’Alessandro, A., Rai, M., and Stark, N. (2025). The intracellular symbiont Wolbachia alters Drosophila development and metabolism to buffer against nutritional stress. PLOS Genet. 21, e1011905. 10.1371/journal.pgen.1011905.

Weeks, A.R., Turelli, M., Harcombe, W.R., Reynolds, K.T., and Hoffmann, A.A. (2007). From Parasite to Mutualist: Rapid Evolution of Wolbachia in Natural Populations of Drosophila. PLOS Biol. 5, e114. 10.1371/journal.pbio.0050114.

Makepeace, B.L., and Gill, A.C. (2016). Wolbachia. In Rickettsiales: Biology, Molecular Biology, Epidemiology, and Vaccine Development, S. Thomas, ed. (Springer International Publishing), pp. 465–512. 10.1007/978-3-319-46859-4_21.

Cornwallis, C.K., van ’t Padje, A., Ellers, J., Klein, M., Jackson, R., Kiers, E.T., West, S.A., and Henry, L.M. (2023). Symbioses shape feeding niches and diversification across insects. Nat. Ecol. Evol. 7, 1022–1044. 10.1038/s41559-023-02058-0.

Hosokawa, T., Koga, R., Kikuchi, Y., Meng, X.-Y., and Fukatsu, T. (2010). Wolbachia as a bacteriocyte-associated nutritional mutualist. Proc. Natl. Acad. Sci. 107, 769–774. 10.1073/pnas.0911476107.

Ju, J.-F., Bing, X.-L., Zhao, D.-S., Guo, Y., Xi, Z., Hoffmann, A.A., Zhang, K.-J., Huang, H.-J., Gong, J.-T., Zhang, X., et al. (2020). *Wolbachia* supplement biotin and riboflavin to enhance reproduction in planthoppers. ISME J. 14, 676–687. 10.1038/s41396-019-0559-9.

Vancaester, E., and Blaxter, M. (2023). Phylogenomic analysis of Wolbachia genomes from the Darwin Tree of Life biodiversity genomics project. PLOS Biol. 21, e3001972. 10.1371/journal.pbio.3001972.

Driscoll, T.P., Verhoeve, V.I., Brockway, C., Shrewsberry, D.L., Plumer, M., Sevdalis, S.E., Beckmann, J.F., Krueger, L.M., Macaluso, K.R., Azad, A.F., et al. (2020). Evolution of Wolbachia mutualism and reproductive parasitism: insight from two novel strains that co-infect cat fleas. PeerJ 8, e10646. 10.7717/peerj.10646.

Nikoh, N., Hosokawa, T., Moriyama, M., Oshima, K., Hattori, M., and Fukatsu, T. (2014). Evolutionary origin of insect– *Wolbachia* nutritional mutualism. Proc. Natl. Acad. Sci. 111, 10257–10262. 10.1073/pnas.1409284111.

Gerth, M., and Bleidorn, C. (2016). Comparative genomics provides a timeframe for Wolbachia evolution and exposes a recent biotin synthesis operon transfer. Nat. Microbiol. 2, 16241. 10.1038/nmicrobiol.2016.241.

Ollerton, J. (2021). Pollinators and Pollination: Nature and Society (Pelagic Publishing Ltd).

Michener, C.D. (2007). The Bees of the World (JHU Press).

Gerth, M., Saeed, A., White, J.A., and Bleidorn, C. (2015). Extensive screen for bacterial endosymbionts reveals taxon-specific distribution patterns among bees (Hymenoptera, Anthophila). FEMS Microbiol. Ecol. 91. 10.1093/femsec/fiv047.

Gerth, M., Röthe, J., and Bleidorn, C. (2013). Tracing horizontal *W olbachia* movements among bees ( A nthophila): a combined approach using multilocus sequence typing data and host phylogeny. Mol. Ecol. 22, 6149–6162. 10.1111/mec.12549.

Stanley, R.G., and Linskens, H.F. (1974). Pollen: Biology Biochemistry Management (Springer Science & Business Media).

Jain, C., Rodriguez-R, L.M., Phillippy, A.M., Konstantinidis, K.T., and Aluru, S. (2018). High throughput ANI analysis of 90K prokaryotic genomes reveals clear species boundaries. Nat. Commun. 9, 5114. 10.1038/s41467-018-07641-9.

Bordenstein, S.R., and Bordenstein, S.R. (2022). Widespread phages of endosymbionts: Phage WO genomics and the proposed taxonomic classification of Symbioviridae. PLOS Genet. 18, e1010227. 10.1371/journal.pgen.1010227.

Vautrin, E., Charles, S., Genieys, S., and Vavre, F. (2007). Evolution and invasion dynamics of multiple infections with *Wolbachia* investigated using matrix based models. J. Theor. Biol. 245, 197–209. 10.1016/j.jtbi.2006.09.035.

Riegler, M., Sidhu, M., Miller, W.J., and O’Neill, S.L. (2005). Evidence for a global Wolbachia replacement in Drosophila melanogaster. Curr. Biol. CB 15, 1428–1433. 10.1016/j.cub.2005.06.069.

Duron, O., and Gottlieb, Y. (2020). Convergence of Nutritional Symbioses in Obligate Blood Feeders. Trends Parasitol. 36, 816–825. 10.1016/j.pt.2020.07.007.

Danforth, B.N., Minckley, R.L., Neff, J.L., and Fawcett, F. (2019). The Solitary Bees: Biology, Evolution, Conservation (Princeton University Press) 10.1515/9780691189321.

Llnskens, H.F., and Jorde, W. (1997). Pollen as food and medicine—A review. Econ. Bot. 51, 78–86. 10.1007/BF02910407.

Morin, A., Rabbat, I., Loranger, Y., Fournier, V., Bergeron, P., MacKell, S., Smale, P., Kerekes, T., Colla, S.R., and Tissier, M.L. (2026). Pol-NIC: An open database on pollen nutrients, imaging, and contaminants. Ecol. Solut. Evid. 7, e70197. 10.1002/2688-8319.70197.

Karalia, S., and Meena, V.K. (2025). Biotin: DNA to diet. J. Nutr. Biochem. 146, 110081. 10.1016/j.jnutbio.2025.110081.

Dharampal, P.S., Hetherington, M.C., and Steffan, S.A. (2020). Microbes make the meal: oligolectic bees require microbes within their host pollen to thrive. Ecol. Entomol. 45, 1418–1427. 10.1111/een.12926.

Steffan, S.A., Dharampal, P.S., Danforth, B.N., Gaines-Day, H.R., Takizawa, Y., and Chikaraishi, Y. (2019). Omnivory in Bees: Elevated Trophic Positions among All Major Bee Families. Am. Nat. 194, 414–421. 10.1086/704281.

Westrich, P. (2019). Die Wildbienen Deutschlands 2. Auflage. (Verlag Eugen Ulmer).

Baldo, L., Dunning Hotopp, J.C., Jolley, K.A., Bordenstein, S.R., Biber, S.A., Choudhury, R.R., Hayashi, C., Maiden, M.C.J., Tettelin, H., and Werren, J.H. (2006). Multilocus Sequence Typing System for the Endosymbiont *Wolbachia pipientis*. Appl. Environ. Microbiol. 72, 7098–7110. 10.1128/AEM.00731-06.

Fernández, M.B., Bleidorn, C., and Calcaterra, L. (2022). Wolbachia Infection in Native Populations of the Invasive Tawny Crazy Ant Nylanderia fulva. Front. Insect Sci. 2, 905803. 10.3389/finsc.2022.905803.

Gansauge, M.-T., and Meyer, M. (2019). A Method for Single-Stranded Ancient DNA Library Preparation. In Ancient DNA Methods in Molecular Biology., B. Shapiro, A. Barlow, P. D. Heintzman, M. Hofreiter, J. L. A. Paijmans, and A. E. R. Soares, eds. (Springer New York), pp. 75–83. 10.1007/978-1-4939-9176-1_9.

Bourlat, S.J., Haenel, Q., Finnman, J., and Leray, M. (2016). Preparation of Amplicon Libraries for Metabarcoding of Marine Eukaryotes Using Illumina MiSeq: The Dual-PCR Method. In Marine Genomics Methods in Molecular Biology., S. J. Bourlat, ed. (Springer New York), pp. 197–207. 10.1007/978-1-4939-3774-5_13.

Chen, S., Zhou, Y., Chen, Y., and Gu, J. (2018). fastp: an ultra-fast all-in-one FASTQ preprocessor. Bioinformatics 34, i884–i890. 10.1093/bioinformatics/bty560.

Li, D., Liu, C.-M., Luo, R., Sadakane, K., and Lam, T.-W. (2015). MEGAHIT: an ultra-fast single-node solution for large and complex metagenomics assembly via succinct de Bruijn graph. Bioinformatics 31, 1674–1676. 10.1093/bioinformatics/btv033.

Challis, R., Richards, E., Rajan, J., Cochrane, G., and Blaxter, M. (2019). BlobToolKit – Interactive quality assessment of genome assemblies. Preprint at bioRxiv, 10.1101/844852 https://doi.org/10.1101/844852.

Li, H. (2018). Minimap2: pairwise alignment for nucleotide sequences. Bioinformatics 34, 3094–3100. 10.1093/bioinformatics/bty191.

Camacho, C., Coulouris, G., Avagyan, V., Ma, N., Papadopoulos, J., Bealer, K., and Madden, T.L. (2009). BLAST+: architecture and applications. BMC Bioinformatics 10, 421. 10.1186/1471-2105-10-421.

Parks, D.H., Imelfort, M., Skennerton, C.T., Hugenholtz, P., and Tyson, G.W. (2015). CheckM: assessing the quality of microbial genomes recovered from isolates, single cells, and metagenomes. Genome Res. 25, 1043–1055. 10.1101/gr.186072.114.

Li, H., Handsaker, B., Wysoker, A., Fennell, T., Ruan, J., Homer, N., Marth, G., Abecasis, G., Durbin, R., and 1000 Genome Project Data Processing Subgroup (2009). The Sequence Alignment/Map format and SAMtools. Bioinformatics 25, 2078–2079. 10.1093/bioinformatics/btp352.

Kolmogorov, M., Yuan, J., Lin, Y., and Pevzner, P.A. (2019). Assembly of long, error-prone reads using repeat graphs. Nat. Biotechnol. 37, 540–546. 10.1038/s41587-019-0072-8.

Wick, R.R., Schultz, M.B., Zobel, J., and Holt, K.E. (2015). Bandage: interactive visualization of de novo genome assemblies. Bioinformatics 31, 3350–3352. 10.1093/bioinformatics/btv383.

Mölder, F., Jablonski, K.P., Letcher, B., Hall, M.B., Tomkins-Tinch, C.H., Sochat, V., Forster, J., Lee, S., Twardziok, S.O., Kanitz, A., et al. (2021). Sustainable data analysis with Snakemake. Preprint at F1000Research, 10.12688/f1000research.29032.1 https://doi.org/10.12688/f1000research.29032.1.

O’Leary, N.A., Cox, E., Holmes, J.B., Anderson, W.R., Falk, R., Hem, V., Tsuchiya, M.T.N., Schuler, G.D., Zhang, X., Torcivia, J., et al. (2024). Exploring and retrieving sequence and metadata for species across the tree of life with NCBI Datasets. Sci. Data 11, 732. 10.1038/s41597-024-03571-y.

Schwengers, O., Jelonek, L., Dieckmann, M.A., Beyvers, S., Blom, J., and Goesmann, A. (2021). Bakta: rapid and standardized annotation of bacterial genomes via alignment-free sequence identification. Microb. Genomics 7, 000685. 10.1099/mgen.0.000685.

Cosentino, S., and Iwasaki, W. (2019). SonicParanoid: fast, accurate and easy orthology inference. Bioinformatics 35, 149–151. 10.1093/bioinformatics/bty631.

Katoh, K., and Standley, D.M. (2013). MAFFT Multiple Sequence Alignment Software Version 7: Improvements in Performance and Usability. Mol. Biol. Evol. 30, 772–780. 10.1093/molbev/mst010.

Minh, B.Q., Schmidt, H.A., Chernomor, O., Schrempf, D., Woodhams, M.D., von Haeseler, A., and Lanfear, R. (2020). IQ-TREE 2: New Models and Efficient Methods for Phylogenetic Inference in the Genomic Era. Mol. Biol. Evol. 37, 1530–1534. 10.1093/molbev/msaa015.

Gerth, M., Saeed, A., White, J.A., and Bleidorn, C. (2015). Extensive screen for bacterial endosymbionts reveals taxon-specific distribution patterns among bees (Hymenoptera, Anthophila). FEMS Microbiol. Ecol. 91, fiv047. 10.1093/femsec/fiv047.

Baldo, L., Dunning Hotopp, J.C., Jolley, K.A., Bordenstein, S.R., Biber, S.A., Choudhury, R.R., Hayashi, C., Maiden, M.C.J., Tettelin, H., and Werren, J.H. (2006). Multilocus sequence typing system for the endosymbiont Wolbachia pipientis. Appl. Environ. Microbiol. 72, 7098–7110. 10.1128/AEM.00731-06.

Camargo, A.P., Roux, S., Schulz, F., Babinski, M., Xu, Y., Hu, B., Chain, P.S.G., Nayfach, S., and Kyrpides, N.C. (2024). Identification of mobile genetic elements with geNomad. Nat. Biotechnol. 42, 1303–1312. 10.1038/s41587-023-01953-y.

R Core Team (2024). R: A Language and Environment for Statistical Computing (R Foundation for Statistical Computing).

Posit team (2025). RStudio: Integrated Development Environment for R (Posit Software, PBC).

Giovannini, M., Gammuto, L., Alonso-Vásquez, T., Gogoleva, N., Bellinzona, G., Potekhin, A., Petroni, G., and Castelli, M. (2026). Facultative mutualism between Paramecium and the intracellular Rickettsiales bacterium Megaera mediated by a horizontally acquired biotin operon. ISME Commun. 6, ycag079. 10.1093/ismeco/ycag079.

Oksanen, J., Simpson, G.L., Blanchet, F.G., Kindt, R., Legendre, P., Minchin, P.R., O’Hara, R.B., Solymos, P., Stevens, M.H.H., Szoecs, E., et al. (2001). vegan: Community Ecology Package. 10.32614/CRAN.package.vegan https://doi.org/10.32614/CRAN.package.vegan.

Baltz, L.M., Gardein, H., Greil, H., Paxton, R.J., and Theodorou, P. (2026). Different Paths, Similar Pressures: Divergent Drivers of Genetic Diversity Despite Convergent Genomic Signatures of Selection in Response to Urban Intensity in Two Oligolectic Bee Species. Mol. Ecol. 35, e70370. 10.1111/mec.70370.

