## Supplementary figures and images for "Rapid spread of nutritional *Wolbachia* symbionts across and within solitary bee species"

### Supplementary Figure 1

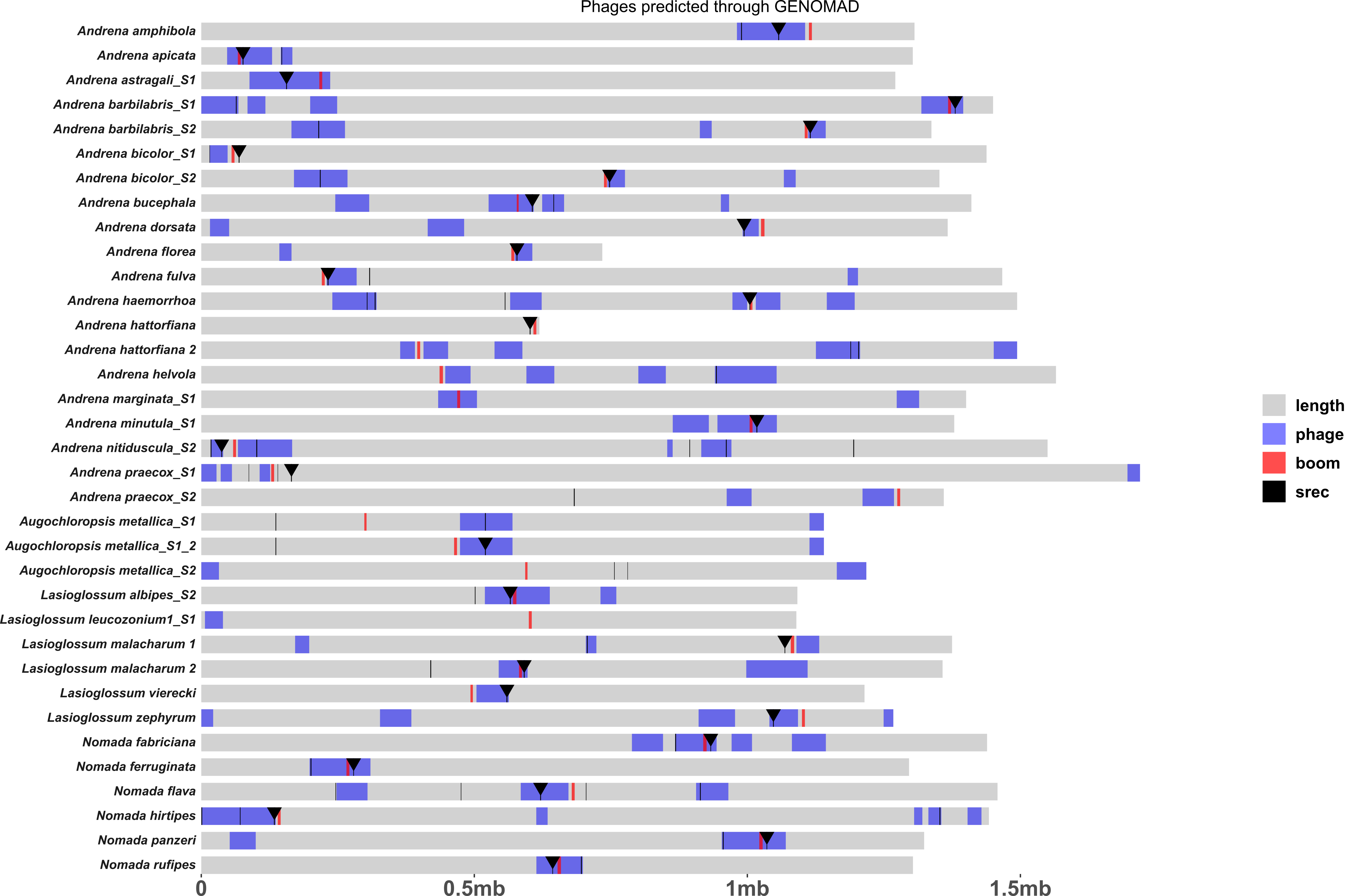

### Supplementary Figure 2

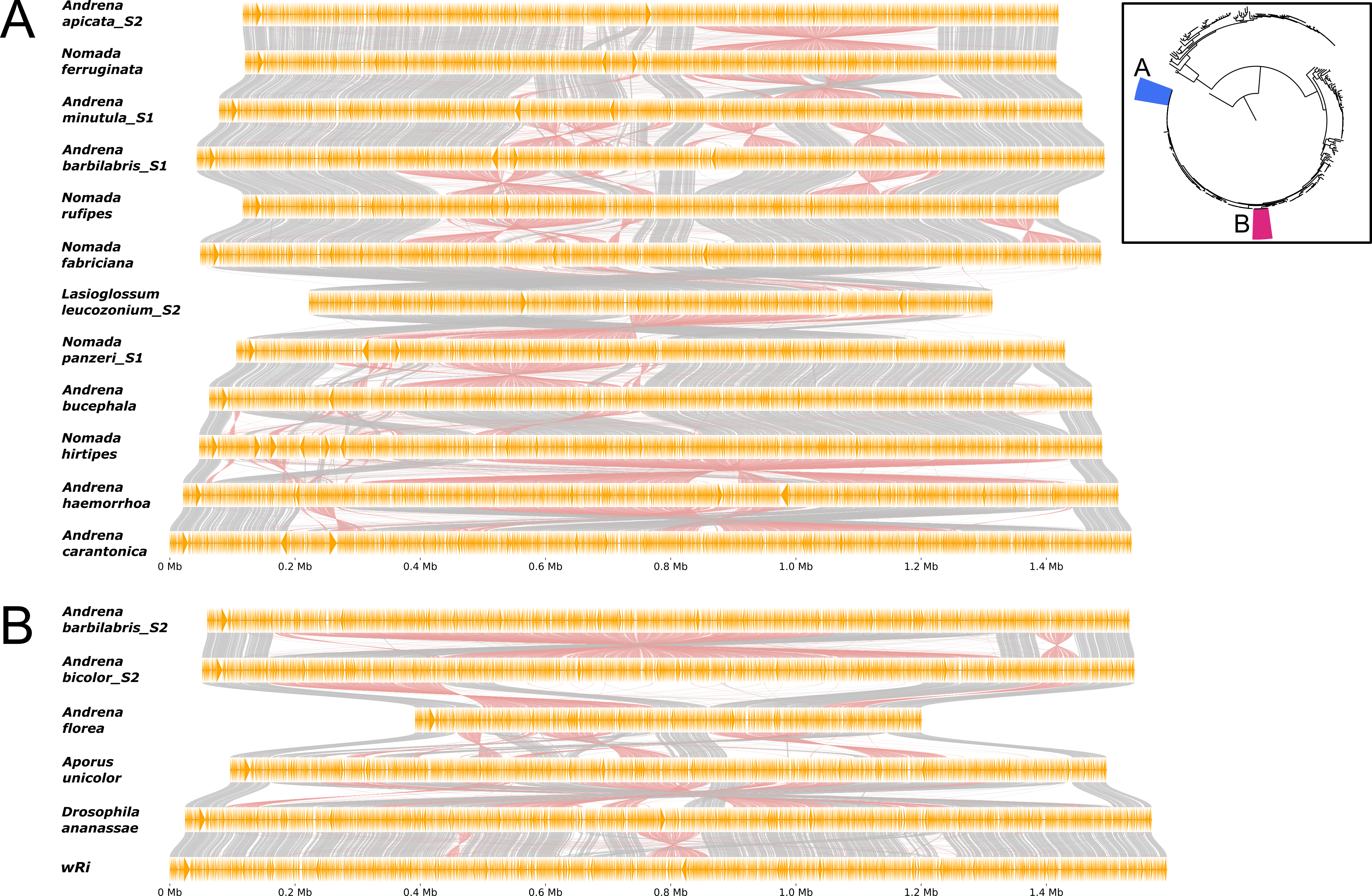

### Supplementary Figure 3

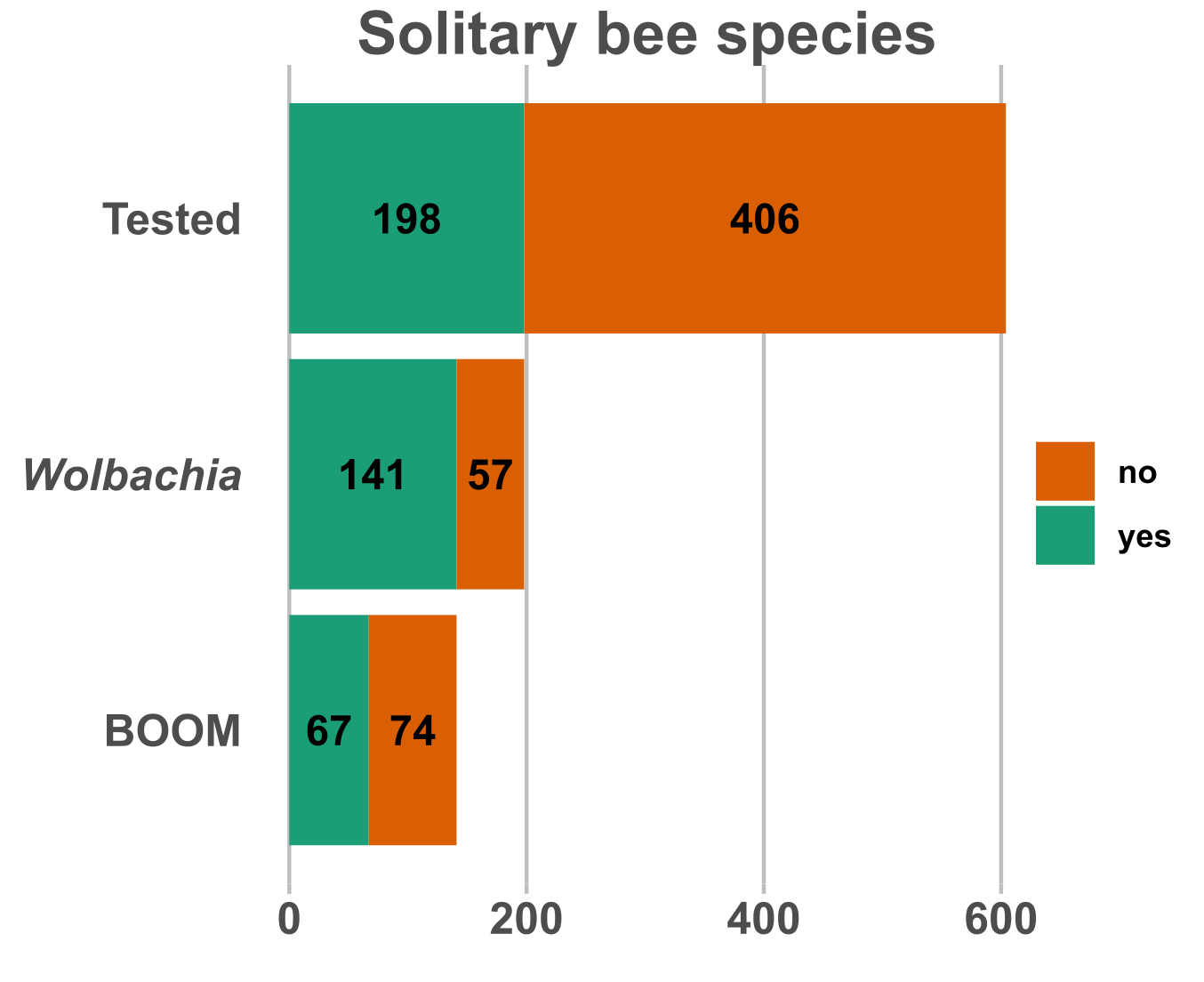
